## Supplementary figures and images for "Structural insight into the stabilization of microtubules by taxanes"

### Movie M5

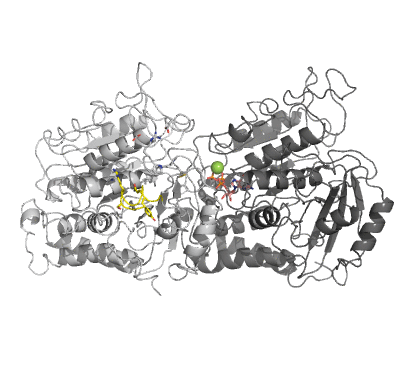

### Movie M6

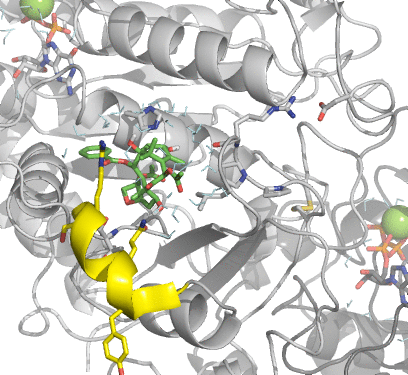

### Movie M7

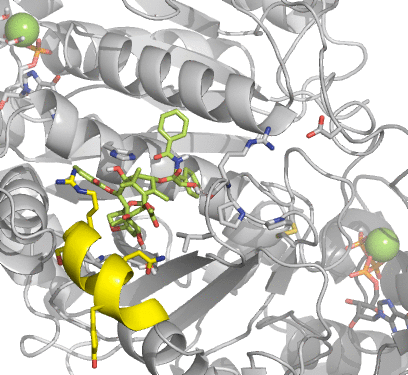
